# Plasmin-Mediated Processing of Clumping Factor A Impacts Bacterial, Aggregation and Abscess Formation during *Staphylococcus aureus* Infection

**DOI:** 10.1101/2025.06.30.662199

**Authors:** Mary B. Turley, Zhicheng Hu, Ross W. Ward, Ed C. Lavelle, Tao Jin, Joan A. Geoghegan

**Author notes:** Corresponding author: Joan A. Geoghegan. Mailing address: Department of Microbes, Infections and Microbiomes, College of Medicine and Health, University of Birmingham.

## Abstract

The interface between pathogenic bacteria and the host plays a critical role during the progression of infection. Bacterial surface components, such as the *Staphylococcus aureus* cell wall-anchored protein clumping factor A (ClfA), interact with host factors. ClfA promotes bacterial aggregation in plasma by binding to fibrinogen. Here, we show that the host serine endopeptidase plasmin cleaves ClfA, disrupting its interaction with fibrinogen and promoting bloodstream survival and virulence in *S. aureus*. We found that plasmin cleaves ClfA between residues Arg214 and Ala215. Cleavage occurs in human serum regardless of whether plasmin becomes activated by host tissue-plasminogen activator or by the activity of the *S. aureus* enzyme staphylokinase. Expressing a plasmin-resistant form of ClfA, and a truncated form that mimics the plasmin-cleaved product on the bacterial cell surface, revealed that following cleavage, ClfA cannot mediate aggregation of *S. aureus* in fibrinogen. Truncated ClfA on the cell surface cannot bind to fibrinogen and promotes the establishment of renal abscesses *in vivo*. Collectively, our findings suggest that activation of plasmin during infection leads to site-specific processing of ClfA, and this not only abrogates fibrinogen binding but also facilitates enhanced *S. aureus* persistence in the host. Thus, we demonstrate that proteolytic processing by plasmin is a mechanism for modulating the host-pathogen interaction during infection.

**Author Summary:** *Staphylococcus aureus* is a major human bacterial pathogen that binds to host factors during infection using its surface adhesins. Here we describe a novel mechanism by which host proteolytic activity modulates bacterial interaction with the plasma protein fibrinogen to influence the course of infection. The surface adhesin clumping factor A (ClfA), binds to fibrinogen, causing bacterial aggregation in the bloodstream. In this study, we show that the host enzyme plasmin cleaves ClfA at a specific site, disrupting its ability to bind fibrinogen. Cleavage eliminates ClfA-mediated aggregation in plasma fibrinogen and enhances *S. aureus* virulence in vivo, increasing the formation of kidney abscesses. Our work therefore demonstrates that specific protease-mediated processing of bacterial adhesins by host factors can influence the progression of infection.

## Introduction

*Staphylococcus aureus* bacteraemia is among the most common and serious bacterial infections affecting the global population (1). *S. aureus* was the leading bacterial cause of death in 135 countries in 2019 (2), and more than 300,000 people die each year as a direct result of *S. aureus* bacteraemia (3). Complications arise when *S. aureus* evades host immune defences and disseminates to organs and tissues giving rise to metastatic infections such as infective endocarditis, septic arthritis, and infections associated with implanted medical devices. These infections increase morbidity and mortality, often requiring prolonged treatment and, in some cases, surgical intervention.

One of the key virulence factors of *S. aureus* is clumping factor A (ClfA), a cell wall-anchored protein that plays a central role during bacteraemia, septic arthritis and infective endocarditis through fibrinogen-dependent and independent mechanisms (4–10). ClfA binds to fibrinogen, a glycoprotein present in blood plasma, promoting the aggregation of *S. aureus* in plasma. However, fibrinogen also functions in host defense by activating neutrophils and macrophages, as well as trapping bacteria in matrices and inhibiting dissemination (11–15). Fibrinogen-deficient mice have reduced ability to clear *S. aureus* from the peritoneal cavity following infection, and disruption of the *S. aureus*-fibrinogen interaction inhibits dissemination from the peritoneum during acute infection (16).

ClfA binds to the extreme C-terminus of the γ chain of fibrinogen. The C-terminal residues of the γ chain bind in a trench formed between the separately folded adjacent immunoglobulin G-like domains N2 and N3 domains by the ‘dock, lock, and latch’ mechanism (17–19). Docking of the peptide induces a conformational change in the flexible extension at the C-terminus of the ClfA N3 domain, which redirects to cover the bound fibrinogen peptide locking it in place. In doing so it forms a β strand which is complementary to a β sheet in the N2 domain. The D domain of fibrinogen establishes further interactions with another site on the surface of the ClfA N3 domain (20). The overall stability of the ClfA-fibrinogen complex is enhanced by shear stress (21). ClfA has also been reported to bind to the host protein factor I, facilitating cleavage of complement opsonin C3b into iC3b to inhibit opsonophagocytosis (10, 22). However, no studies have yet detailed the specific mechanism or binding site involved.

There is growing evidence that host-derived proteases alter the *S. aureus* cell surface, potentially impacting its interactions with the host during infection (23–25). The host serine protease plasmin is particularly relevant in the context of infection due to its activation at sites of tissue damage and inflammation (26, 27). Notably, *S. aureus* produces staphylokinase (Sak), a secreted protein that activates host plasminogen to its proteolytically active form, plasmin (28). This interaction suggests a potential mechanism by which *S. aureus* may exploit the host fibrinolytic system to remodel its own cell surface. This study set out to determine if ClfA is susceptible to plasmin and whether plasmin can modulate the biological activity of ClfA.

## Materials and Methods

### Reagents, Strains, Media, and Growth Conditions

All strains and plasmids used in this study are listed in Table 1. If not specified otherwise, *S. aureus* strains were cultured on tryptic soy agar (TSA) plates for isolation of individual colonies or for liquid cultures in tryptic soy broth (TSB) (Difco) at 37°C with shaking (200 rpm). *Escherichia coli* strains were grown at 37°C on lysogeny Agar (LA) or in lysogeny Broth (LB) with shaking (200 rpm) for liquid cultures. Where appropriate, antibiotics were incorporated into media; chloramphenicol (Cm,10 μg/mL), erythromycin (Em, 10 μg/mL), ampicillin (Amp, 100 μg/mL); anhydrotetracycline (Atc, 1 ng/mL) (Sigma).

### Plasmin treatment

Bacteria were grown overnight for 16-20 h, then pelleted by centrifugation (7 min, 8000 x *g*), washed with an equal volume of PBS once and pelleted again. Samples were then resuspended in 1 mL PBS and adjusted to the desired OD_600_ for the assay. Human plasmin (Sigma) was added to the sample to a final concentration of 0.075 U/mL and incubated at 37°C for 1 h in a water bath. Samples were then washed with an equal volume of PBS three times before a final reconstitution to original volume. The protease inhibitor alpha-2-macroglobulin (α2M, Sigma) was added where indicated at a concentration of 10 times that of plasmin used and preincubated with plasmin for 20 minutes to inactivate the protease. Recombinant proteins were typically diluted to a concentration of 5 μM in sterile PBS to which plasmin was added to a final concentration of 0.075 U/mL and incubated at 37°C for 1 h in a water bath.

### Preparation of human and murine blood and serum

Fresh blood samples were drawn from healthy human volunteers, into serum clot activator vacuette tubes (Greiner) or EDTA-containing vacuettes (Greiner). Serum was collected by inverting the serum clot activator tubes, then incubating at room temperature for 15 min before centrifuging at 1,500 x g for 16 min. To collect murine serum, whole blood was collected from female C57BL6/JCR mice by tail bleed and was allowed to clot overnight at 4°C. Serum was then harvested from clotted blood by centrifugation at 1100 x *g* for 11 min. To catalyse the activation of plasminogen to plasmin in human serum, 40% serum was incubated with tissue plasminogen activator (DiaPharma, 25 µg/mL) or *S. aureus* supernatant for 20 min at 37°C before being used in the treatment of recombinant proteins as outlined above.

### SDS-PAGE and Western immunoblotting

Cell wall extractions were carried out as described by O’Halloran *et al*., 2015 (29). Briefly, bacteria grown to exponential phase (OD_600_ = 0.35 - 0.5) or stationary phase (overnight culture) were pelleted and washed with PBS, then adjusted to an OD_600_ = 10. Bacteria were then incubated with or without plasmin (0.075 U/mL) in a 37°C waterbath for 1 h before being centrifuged and resuspended in 250 μL extraction solution (30% w/v Raffinose, 50 mM Tris-HCL, 20 mM MgCl2, pH 7.5, 50 µl/ml complete protease inhibitor [Roche], lysostaphin 10 U/mL) for 8 min at 37°C. Protoplasts were removed by centrifugation at 15000 x *g* for 5 minutes.

Protein samples were separated by polyacrylamide gel electrophoresis (7.5%, 10% or 12.5%). Gels were either stained using Instant Blue (Expedeon) or Gel code (ThermoFisher Scientific) or transferred to nitrocellulose (GE Healthcare) or polyvinylidene difluoride (PVDF, Roche) filters for western immunoblotting.

Filters were blocked with 10% (w/v) skimmed milk (Marvel) and then probed with polyclonal anti-ClfA N1N2N3 IgG (1:1000, (30)) followed by horseradish peroxidase (HRP)-conjugated protein A (1:500, Sigma), 7E8 IgG (1:1000 (31)) followed by goat anti-mouse IgG-HRP (1:2500, Dako) or alternatively with Strep-tag® II IgG-HRP (1:2500, Novagen). Filters were developed using LumiGLO Reagent and peroxide detection solution (Cell signalling Technology) and imaged using the ImageQuant TL (GE).

### DNA manipulation

#### Isolation of DNA

Genomic DNA was isolated using the PureElute Bacterial genomic DNA Purification kit (Edge Biosystems), with the addition of lysostaphin (100 µg/ml) to the cell lysis buffer to digest *S. aureus* cell wall peptidoglycan to improve DNA extraction. Plasmid DNA was isolated using Wizard® Plus SVMiniprep kit (Promega) or Qiagen® Plasmid plus Midi-kit.

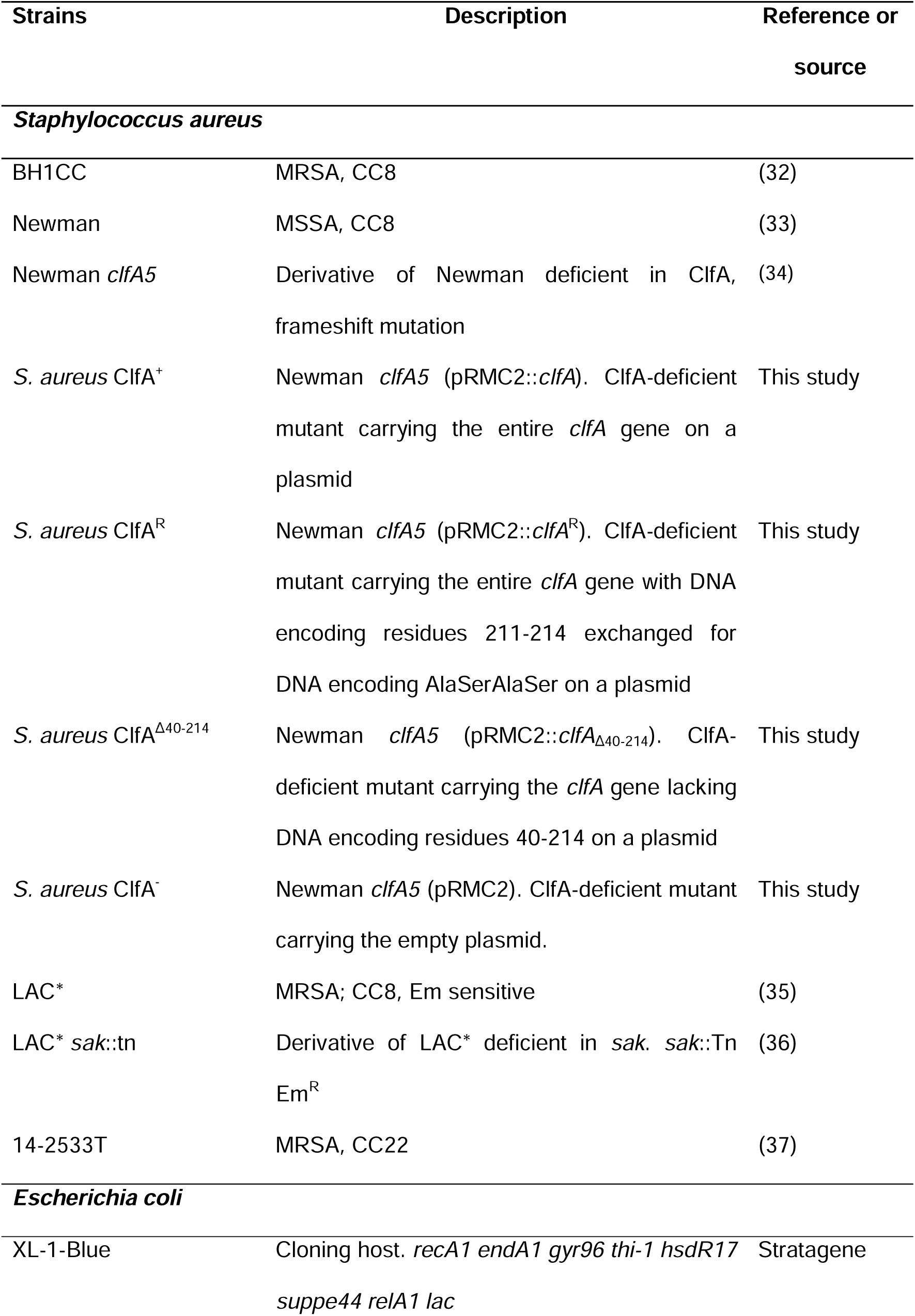
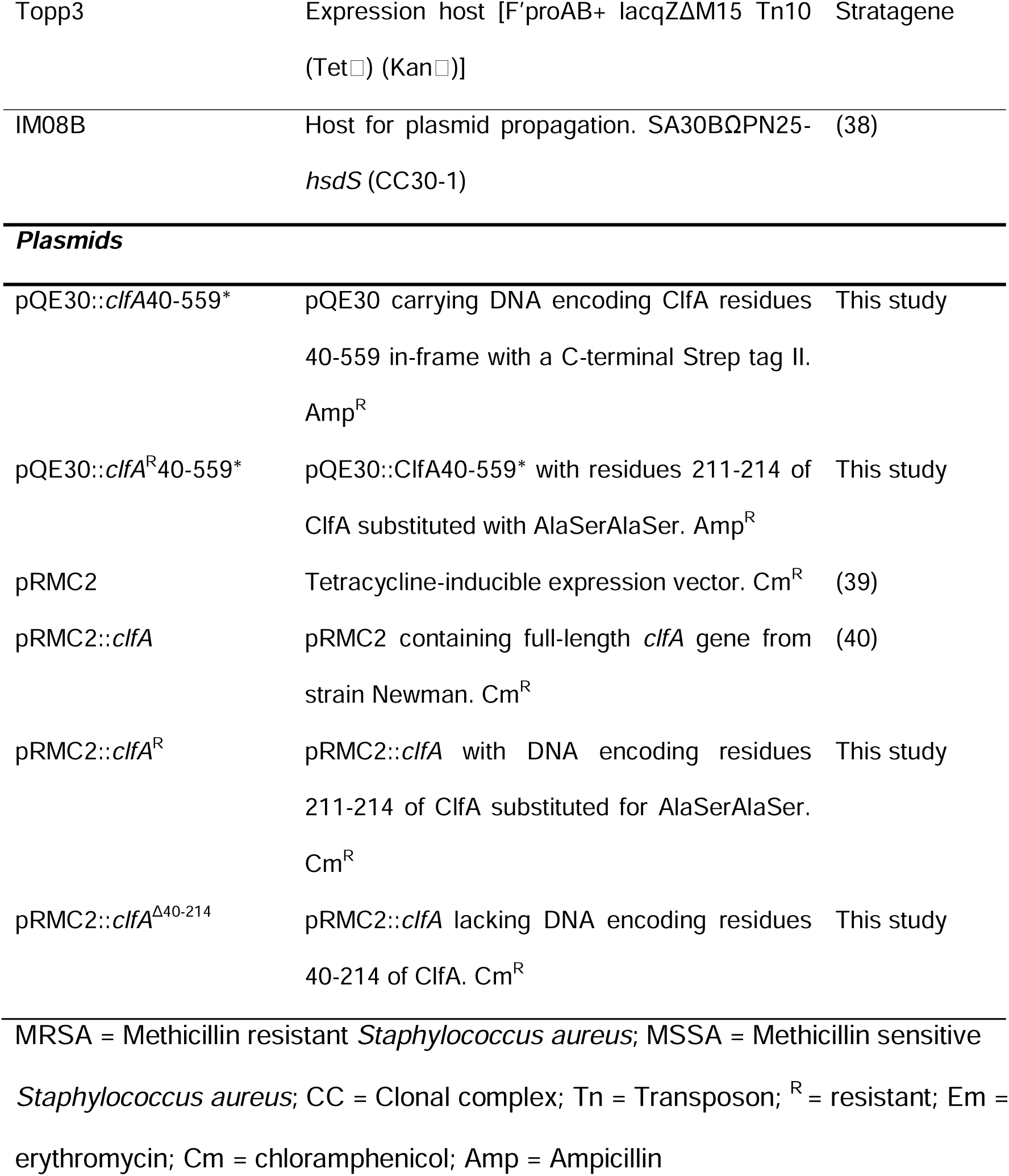

#### Plasmid construction

Plasmids were constructed by inverse PCR or via sequence and ligase independent cloning (51). Plasmids pQE30::ClfA^R^_40-559*_ and pRMC2::ClfA^R^ were constructed by sequence and ligase independent cloning (41) where vectors were amplified using Velocity polymerase and the insert was amplified from a region of DNA commercially synthesised as a gBlock (IDT). Primers used in this study are listed in **Table S1**. Plasmids pQE30::ClfA_40-559*_ and pRMC2::ClfA^Δ40-214^ were generated by inverse PCR. Inverse PCR products were treated with DpnI and blunt-end ligation was carried out using the LigaFast® Rapid RNA ligation system (Promega). Chemically competent *E. coli* (42) XL-1 Blue was transformed with the ligation products. Plasmids were then isolated and used to transform *E. coli* Topp3 made chemically competent for protein purification or *E. coli* IM08B for transformation of *S. aureus* made electrocompetent as described previously (43). Colony PCRs were carried out to screen for transformants using Phire Hot Start DNA Polymerase (Sigma) as per the manufacturer’s instructions.

### Expression and purification of recombinant proteins

Briefly, *E. coli* Topp3 carrying plasmids was grown to an OD_600_ = 0.4 - 0.8, before isopropyl β-D-1-thiogalactopyranoside (IPTG, 1 mM) was added to induce protein expression. Growth was continued for 3 h before cell sedimentation was carried out. The pellet was resuspended in 25 mL Imidazole Buffer (40 mM Tris-HCL, 1 M NaCl, 0.1 M imidazole, pH 7.9) with an EDTA-free protease inhibitor. Bacteria were then lysed using a French Press at 1500 psi before centrifuging the lysates at 18,000 x *g* for 20 mins at 4°C. DNase (1 mg/mL) was added to the supernatant before filtration through a 0.45 μm filter (Whatman). Proteins were purified with an N-terminal 6xhexa-histidine (His) affinity tag and a C-terminal Strep-tag II affinity tag for purification via nickel affinity chromatography (5 mL HiTrap affinity column; GE Healthcare) and Strep-Tactin affinity chromatography (Strep-Spin Protein Miniprep kit; Zymo Research) respectively. Protein concentrations were determined by measuring the absorbance at 280 nm using a Nanodrop spectrophotometer and calculations according to Beer-Lambert Law.

### N-terminal sequencing

Samples sent for N-terminal sequencing were first incubated with plasmin (0.075 U/mL) as outlined above prior to purification of the digested product by Strep-Tactin affinity chromatography. Purified samples were then subjected to SDS-PAGE, transferred to PVDF membrane and stained with Ponceau S (ThermoScientific). N-terminal sequencing was performed by Edman degradation at Alta Bioscience, Redditch, UK.

### Synthetic peptide cleavage

Synthetic peptides used in this study were commercially synthesised by AltaBioscience (Redditch, UK). Peptide size and purity was confirmed by mass spectrometry at AltaBioscience. All peptides were dissolved in molecular grade water to a concentration of 150 μM. As the peptides were 5/6-FAM labelled they were protected from light when in use and stored at −20°C. Synthetic peptides were diluted to a concentration of 37.5 μM in molecular grade water, then incubated with human plasmin (0.075 U/mL) at 37°C for 30 min in a water bath. Reaction products were separated by agarose gel electrophoresis (4% w/v) and gel images were captured under UV light.

#### Bacterial clumping in fibrinogen

Bacteria were grown to stationary phase, then pelleted by centrifugation (8000 x *g,* 7 min). Cells were washed with PBS and then adjusted to OD_600_ of 1.5 in 1 mL PBS. Purified human fibrinogen depleted of plasminogen, von Willebrand Factor and fibronectin (Enzyme Research Labs) or murine fibrinogen (Innovative Research) was added to the tube to a final concentration of 18.4 μg/mL and left to stand at 37°C. After incubation for 1 h 45 min, the top 0.5 mL was removed from the tube, diluted 1:2 in PBS for OD_600_ measurements (Genesys 10 UV, Thermo Spectronic) and calculated as a percentage of the starting OD_600_ where (100/T0_OD_600_)*T105_OD_600_, where T0 is the starting OD_600_ and T105 is the OD_600_ after 1 h 45 min. For each *S. aureus* strain, there was a fibrinogen-deficient control. In cases where plasmin treated bacteria were included, on adjustment to an OD_600_ = 1.5 the bacteria were incubated with plasmin (0.075 U/mL) for 1 h at 37°C. Following three washes in PBS, the bacteria were resuspended to the original volume in BHI and made up to a final volume of 1 mL with PBS. Fibrinogen (18.4 ug/mL) was then added, and the protocol followed as outlined above. In each case for plasmin treatment, a no plasmin control for each strain was included.

### *Ex vivo* whole blood survival assay

The quantification of survival of *S. aureus* in human blood was performed according to the protocol previously described by Zapotoczna *et al*., 2018 (37). Briefly, *S. aureus* was grown overnight in RPMI then adjusted to 2 × 10^4^ cfu/mL. 25 μL was added to 475 μL of freshly drawn EDTA-treated blood obtained from human volunteers. Tubes were inverted and 100 μL of each sample was immediately added to 900 μL ice-cold endotoxin free water and serial dilutions were made in water for plating on agar to establish the CFU/mL of the sample at T0. The tubes were then incubated at 37°C on a 360° rotation wheel and after 3 h (T3), serial dilutions were made and bacteria plated on agar to establish the number of CFU per mL. In parallel, bacteria were incubated in serum derived from the same donor. Percentage survival was calculated using the following formula (T3 CFU/mL)/(T0 cfu/mL)*100.

### Mouse model of *S. aureus* bloodstream infection

Female NMRI, aged 9-10 weeks, were obtained from Envigo (Venray, Netherlands) and housed at the animal facility at the University of Gothenburg. Mice were maintained under standard temperature and light conditions, with unrestricted access to laboratory chow and water. *S. aureus* strains were cultured for 24 hours on horse blood agar plates and then preserved according to established protocols (44). Before each experiment, the bacterial solutions were thawed, washed, and adjusted to the desired concentration. To investigate the clinical relevance of ClfA plasmin cleavage, we used a well-established *S. aureus* sepsis mouse model (45). NMRI mice (10 per group) were intravenously injected via the tail vein with 200 μL of PBS containing 75 ng/mL anhydrotetracycline and the respective *S. aureus* strain Newman *clfA5* expressing pRMC2 variants: ClfA^+^, ClfA^R^ or ClfA^Δ40-214^ (6 × 10^6^Dcolony-forming units [CFU]/mouse). Expression of ClfA was induced via an intraperitoneal injection of inducer anhydrotetracycline (5 µg/mouse) on Day 0, 1 h before infection and 1h post-infection. The mice were monitored individually two times daily by an observer (Z.H). The blood samples were collected on both day 3 and day 10 for subsequent Interleukin 6 (IL-6) level analysis. Surviving mice were sacrificed on day 10, the blood, kidneys and livers were aseptically collected.

### Quantification of bacterial load in murine kidneys and livers

Kidneys and livers were collected aseptically, and kidney abscess scores were assessed by two investigators (T.J. and Z. H.) in a blinded manner (46). The scoring system used ranged from 0 to 3 (0 indicates healthy kidneys; 1, 1–2 small abscesses in the kidneys without structure changes; 2, >2 abscesses, but <75% kidney tissue involved; and 3, large amounts of abscess with >75% kidney tissue involved). The kidneys and livers were homogenized and plated on horse blood agar plates to quantify the CFU counts.

### Quantification of interleukin 6 using enzyme-linked immunosorbent assay

The levels of Interleukin 6 (IL 6) in the blood of NMRI mice infected with *S. aureus* were measured on day 3 and 10 post-infection using a DuoSet ELISA kit (R&D Systems, Abingdon, UK) according to the manufacturer’s instructions.

### Ethical Approval

Our research complies with all relevant ethical regulations. Informed consent was obtained from all adult participants. Ethical approval for the collection and use of human blood samples from adult volunteers was obtained from the Trinity College Dublin Faculty of Health Sciences Ethics Committee, Ireland (reference 2020111) and through the Human Biomaterials Resource Centre, University of Birmingham, United Kingdom (reference 25/NW/0013). Personal information such as sex and age were not recorded. All participants were over the age of 18 and provided written informed consent. Participants did not receive compensation for their participation. For animal studies conducted in Trinity College Dublin, C57BL/6 mice were bred in-house in the Comparative Medicine Unit, as per university and The Health Products Regulatory Authority (HPRA) welfare guidelines. All animal studies were approved by the TCD Animal Research Ethics Committee (Ethical Approval Number 091210) and HPRA license AE191364/P079. For mouse work, the study was approved by the Gothenburg Ethics Committee of Animal Research (Ethical Approval Number 5.8.18-02443/2021), and all procedures adhered to established guidelines for animal experimentation.

### Statistical analysis

Statistical analysis was performed using GraphPad Prism (GraphPad Software, Inc, USA; version 8.0.2.263 and version 10). Unless stated otherwise, the data presented in this study show mean ± standard deviations of at least three independent experiments. Statistically significant differences were calculated by using methods as indicated in the figure legends. P values of less than 0.05 were considered significant with *, **, *** and **** representing P values of <0.05, <0.01, <0.001 and <0.0001 respectively. P values of >0.05 represents non-significant values (ns). For experiments with blood or serum, samples from different donors were used for each of the biological replicates.

## Results

### ClfA is cleaved by human plasmin

To determine if cell wall-anchored ClfA was susceptible to plasmin-mediated proteolytic processing in *S. aureus*, bacteria were grown to stationary phase, washed and incubated with plasmin. Cell wall-associated proteins were solubilised from the cell wall using lysostaphin during protoplast formation. Probing cell wall extracts of wild-type *S. aureus* Newman with anti-ClfA IgG in a western blot revealed a ca. 170 kDa immunoreactive band corresponding to full length ClfA (**Fig. 1A**). Incubation with plasmin resulted in the appearance of a ca. 130 kDa band in addition to the ca. 170 kDa band indicating that cell-wall anchored ClfA had been cleaved by plasmin. The broad-spectrum protease inhibitor alpha-2-macroglobulin prevented the activity of plasmin as only the band corresponding to full-length ClfA was detected (**Fig. 1A**). As a control, cell wall extracts were prepared from a mutant deficient in ClfA (Newman *clfA5*). The absence of immunoreactive bands confirmed the specificity of the anti-ClfA IgG (**Fig. 1A**). These results suggested that ClfA on the surface of *S. aureus* Newman can be cleaved by purified plasmin.

**FIG. 1.**
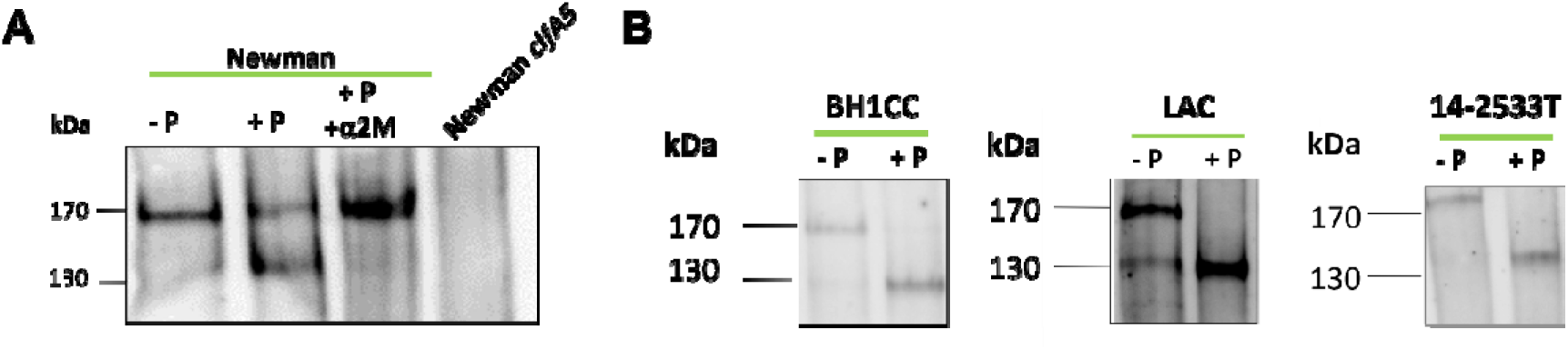
Cell wall-associated ClfA is cleaved by human plasmin. Stationary phase cultures of *S. aureus* strains Newman, Newman *clfA5*, BH1CC, LAC and 14-2533T were adjusted to an OD_600_ = 10 and incubated at 37°C for 1 h with and without plasmin (± P, 0.075 U/mL) and alpha-2 macroglobulin (+ α2M, 1 mg/mL) before cell wall proteins were extracted and separated by SDS-PAGE and probed with anti-ClfA A domain polyclonal IgG (1:1000) in a western blot. Bound primary antibodies was detected with protein A peroxidase (1:500). Blots presented are representative of two independent experiments.

To explore plasmin cleavage further, three methicillin-resistant *S. aureus* (MRSA) isolates were incubated with plasmin and the integrity of ClfA in cell wall extracts was examined. For each of the isolates, ClfA was cleaved by plasmin as indicated by the disappearance of the ca. 170 kDa band corresponding to full-length ClfA and the appearance of a ca. 130 kDa immunoreactive band following plasmin treatment (**Fig. 1B**). A weak immunoreactive band migrating around 130 kDa was detected in cell wall extracts from LAC that had not been incubated with plasmin. This was likely due to the activity of the *S. aureus* extracellular protease aureolysin that cleaves at the junction of N2 and N3 of ClfA (47). In summary, these findings demonstrate that plasmin cleaves ClfA on the surface of *S. aureus*.

Recognition by an antibody specific for ClfA domains N1N2N3 (**Fig. 1**) suggested that ClfA undergoes cleavage towards the N-terminus, leaving a portion of the N1N2N3 domains intact and anchored to the cell wall. To examine this further, a recombinant protein corresponding to the N1N2N3 domains of ClfA with an N-terminal hexahistidine tag and a C-terminal Strep-tagII was purified from *E. coli* (ClfA_40-559*_). ClfA_40-559*_ migrates aberrantly with an apparent molecular mass of ca. 95 kDa (**Fig. 2A**) due to the presence of the elongated unfolded N1 region (48). Incubation of ClfA_40-559*_ with plasmin resulted in the conversion of the ca. 95 kDa band to a single product of ca. 43 kDa. The remainder of the protein could not be detected (**FIG. 2A, B**) suggesting that it was digested into smaller fragments. ClfA_40-559*_ remained mostly intact when the protease activity of plasmin was inhibited by alpha-2-macroglobulin (**FIG. 2A, B**) confirming that cleavage of ClfA_40-559*_ was due to the proteolytic activity of plasmin.

**FIG. 2.**
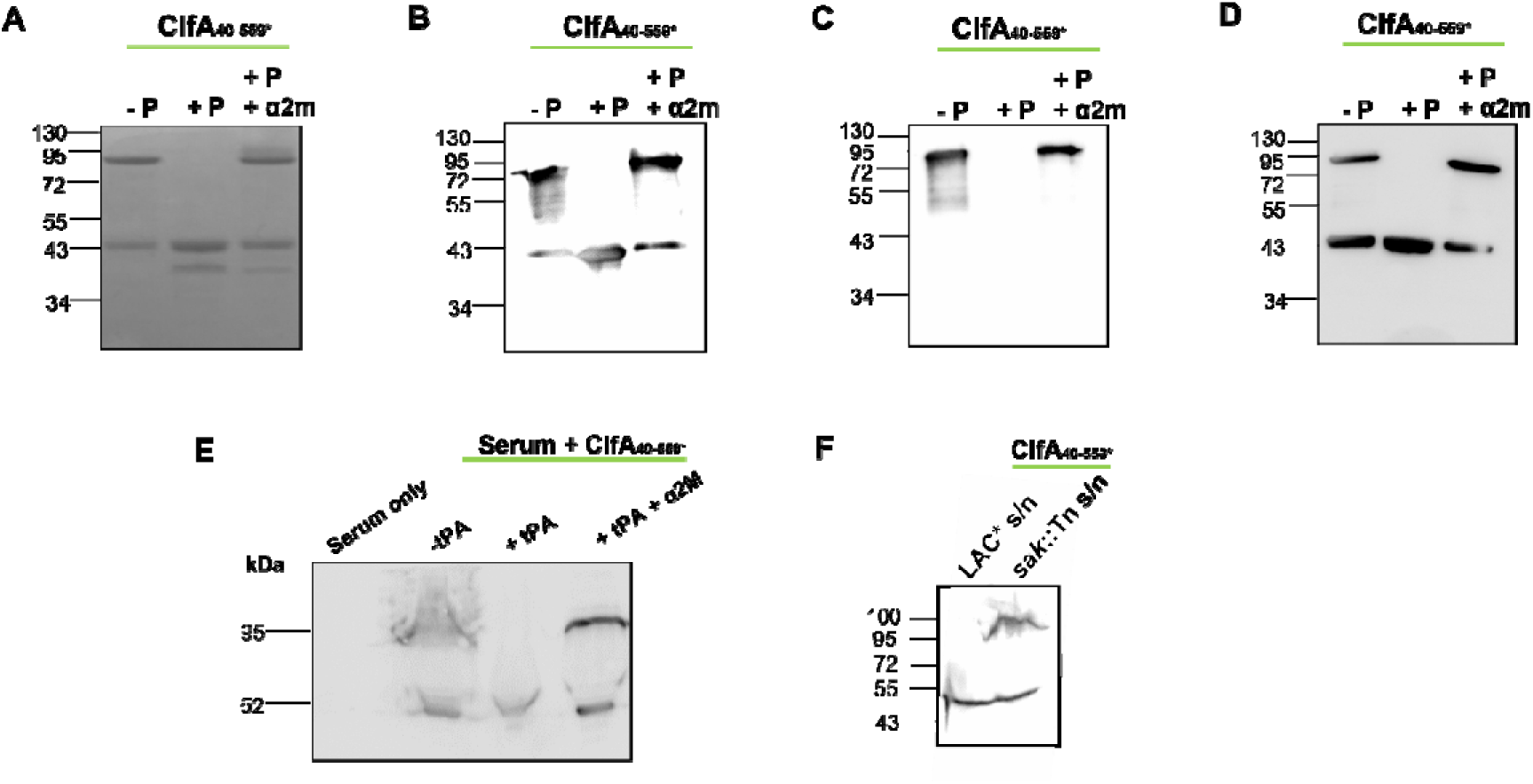
The ClfA N1N2N3 domains are cleaved by human plasmin. ClfA_40-559*_ (5 μM) was incubated in PBS with or without added human plasmin (± P, 0.07 U/mL) and alpha-2 macroglobulin (+ α2M, 1 mg/mL), where indicated, at 37°C for 1 h (A-D). Alternatively, ClfA_40-559*_ (5 μM) was incubated in human serum (40%) that was pretreated with either E) tissue plasminogen activator (± tPA, 25 μg/mL) or F) concentrated supernatants (s/n) from *S. aureus* strain LAC* or its *sak-*deficient mutant (*sak*::Tn) for 20 min at 37°C. Serum only controls did not include ClfA_40-559*_. Samples were separated by SDS-PAGE and either stained with SimplyBlue Coomassie stain (A) or probed in a western immunoblot with anti-ClfA IgG (1:1000; B, E, F), anti-His 7E8 IgG (1:1000; C) or anti-Strep Tag II IgG-HRP (1:2500; D). Bound anti-ClfA IgG was detected with protein A peroxidase (1:500) and bound anti-His IgG was detected with HRP-conjugated rabbit anti-mouse IgG (1:2500). Blots represent two independent experiments.

To determine if the single band remaining following plasmin treatment retained the N-terminal and/or C-terminal tags, the same samples were probed with anti-His IgG to detect the N-terminal hexa-histidine tag (**FIG. 2C**) or anti-Strep tagII IgG to detect the C-terminal strep tag (**Fig. 2D**). While a band corresponding to full-length ClfA_40-559*_ was detected in samples without plasmin or when plasmin was inhibited with alpha-2-macroglobulin, using anti-His IgG, no band was detected after the addition of plasmin (**FIG. 2C**). These data suggest that the hexa-histidine tag, and therefore the N-terminal region of the recombinant protein, is removed by plasmin cleavage. In contrast, the plasmin-cleaved product reacted with anti-Strep tag II IgG (**FIG. 2D**) indicating that the C-terminus of the protein was intact. Other bands detected in lanes 1 and 3 (**FIG. 2B, D**) are likely to be ClfA breakdown product(s) that were already in the sample or formed during the incubation process.

To elucidate if ClfA can be cleaved by activated plasmin in human serum, ClfA_40-559*_ was incubated in serum with or without added tissue plasminogen activator (tPA) which catalyses activation of the zymogen plasminogen (**Fig. 2E**) or with concentrated culture supernatants containing the secreted *S. aureus* plasminogen activating-enzyme Sak (**Fig. 2F**). Full length ClfA_40-559*_ was detected following incubation in serum, but only a band corresponding to cleaved ClfA was detected when plasmin was activated (**Fig. 2E**). The addition of alpha-2-macroglobulin protected full-length ClfA from cleavage. The presence of some truncated protein in the absence of tPA or when plasmin activity was inhibited by alpha-2-macroglobulin indicates that this species was present in the protein preparation at the onset of the experiment or developed independently of plasmin activation. No immunoreactive bands were detected in serum only controls confirming the specificity of the polyclonal antibody for ClfA. Full length ClfA_40-559*_ was detected following incubation in serum treated with concentrated supernatants from an *S. aureus* mutant deficient in *sak* (LAC* *sak*::Tn) but not when ClfA was incubated with supernatants from wild-type LAC* (**Fig. 2F)**, indicating that plasmin was activated by Sak and cleaved ClfA (**Fig. 2F**). Taken together, these findings indicate that purified human plasmin, or plasmin activated by the host or bacterium during infection, is capable of cleaving ClfA.

### Plasmin cleaves ClfA between residues 214 and 215

To identify the plasmin cleavage site, ClfA_40-559*_ was incubated with plasmin and the cleaved protein was purified by Strep-Tactin affinity chromatography. The 7 most N-terminal amino-acids of the cleaved product were identified using Edman degradation, revealing that plasmin cleaves the peptide bond after Arg_214,_ towards the end of the N1 domain (**Fig. 3A**). To verify the plasmin cleavage site experimentally, an 11-mer fluorescein labelled peptide comprising ClfA residues 211-221, was incubated with plasmin and analysed by agarose gel electrophoresis. The 11-mer peptide was chosen based on the evidence from N-terminal sequencing that plasmin would cleave between residues 214 and 215 resulting in a clear size shift from a fluorescein-labelled 11-mer peptide to a fluorescein-labelled 4-mer. In agreement with this, the 11-mer peptide (WT) underwent complete conversion to a smaller truncate following plasmin treatment (**Fig. 3B**). Next a series of peptides where amino-acids 211-216 were individually substituted with serine were tested for susceptibility to plasmin cleavage (**Fig. 3B**). Only the R214S substitution affected cleavage with this peptide undergoing partial truncation following plasmin treatment underscoring the importance of residue Arg214 for plasmin recognition and/or cleavage. A scrambled version of the 11-mer peptide (SD) and peptide where the entire plasmin-cleavage site and surrounding residues were substituted (amino-acids 211-214, WTD) remained intact post-incubation with plasmin. Similarly, recombinant ClfA_40-559*_ harbouring amino-acid substitutions of residues 211-214 (ClfA^R^_40-559*_) was resistant to plasmin cleavage (**Fig. 3C**). Overall, these data indicate that substitution of residues 211-214 prevents the cleavage of ClfA by plasmin and that R214 is particularly important for optimal cleavage by plasmin. Failure of plasmin to cleave the scrambled peptide indicates that the order of the residues is critical for plasmin to cleave this sequence.

**FIG. 3.**
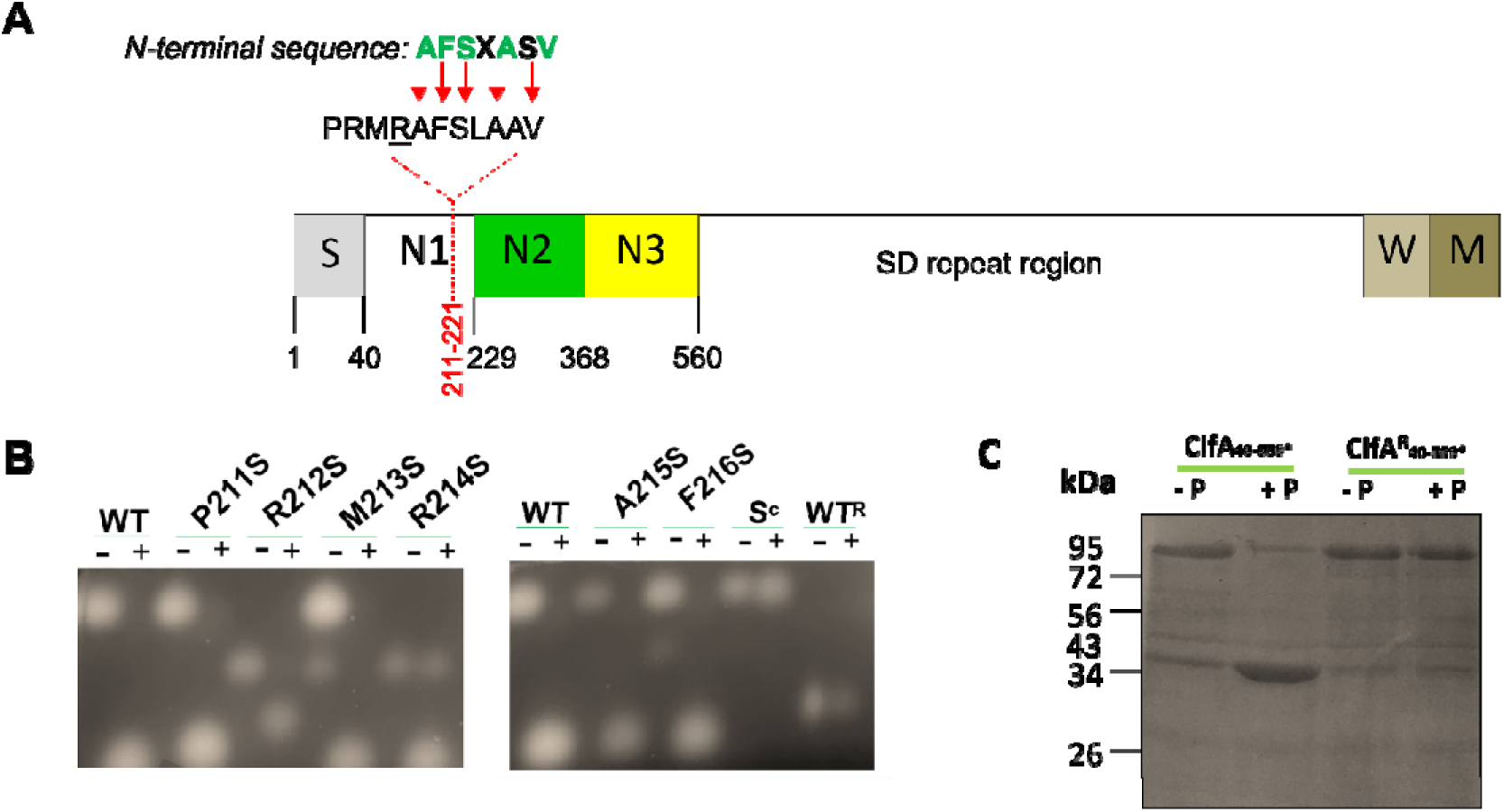
Plasmin cleaves ClfA between residues 214 and 215. A) Domain organisation of ClfA. The amino acid coordinates of the signal peptide (S) and N1N2N3 domains are indicated. The SD repeat region and wall (W) and membranes-spanning (M) regions are shown. The 7 most N-terminal amino-acids of the plasmin-cleaved product identified using Edman degradation are indicated. Sequence identity is indicated in green with red arrows highlighting matches to the ClfA sequence. Arg214 predicted to be cleaved by plasmin is underlined (<u>R</u>). B) A fluorescently-labelled 11-mer peptide (WT) encompassing the region around the proposed cleavage site at Arg214 (residues 211-221) or variants P211S, R212S, M213S, R214S, A215S and R216S harbouring single amino-acid substitutions with serine (37.5 μM) were incubated in the absence (−) or presence (+) of plasmin (0.075 U/mL) for 30 min at 37°C. Peptide S^c^ was a scrambled version of the WT peptide and WT^R^ a peptide with four substitutions (P211A, R212S, M213A, R214S). Samples were analysed by agarose (4% w/v) gel electrophoresis. Gels represent three independent experiments. C) Recombinant ClfA_40-559*_ and ClfAlll_40-559*_ (5 μM) were incubated with or without plasmin (± P, 0.075 U/mL) at 37°C for 1 h and analysed by SDS-PAGE. Gels were stained with Instant Blue Coomassie stain. The gel image presented is representative of two independent experiments.

### Plasmin cleavage disrupts the fibrinogen binding activity of ClfA

ClfA binds fibrinogen in human plasma which facilitates clumping of *S. aureus* by way of the formation of fibrinogen bridges between bacteria (4, 49). The ability of ClfA to interact with fibrinogen was studied by incubating *S. aureus* in a solution of fibrinogen and monitoring the optical density at 600 nm of the suspension as the bacteria clump. A suspension of *S. aureus* expressing full-length ClfA (*S. aureus* ClfA^+^) treated with plasmin to promote processing of ClfA prior to incubation with fibrinogen, underwent significantly less clumping than untreated bacteria (**Fig. 4A**). Conversely, bacteria expressing ClfA carrying amino-acid substitutions in residues 211-214 that render the protein resistant to plasmin cleavage (*S. aureus* ClfA^R^) did not change clumping phenotype following plasmin treatment (**Fig. 4B**). Together these results demonstrate that the cleavage of ClfA by plasmin reduces the protein’s ability to interact with soluble fibrinogen and promote clumping.

**FIG. 4.**
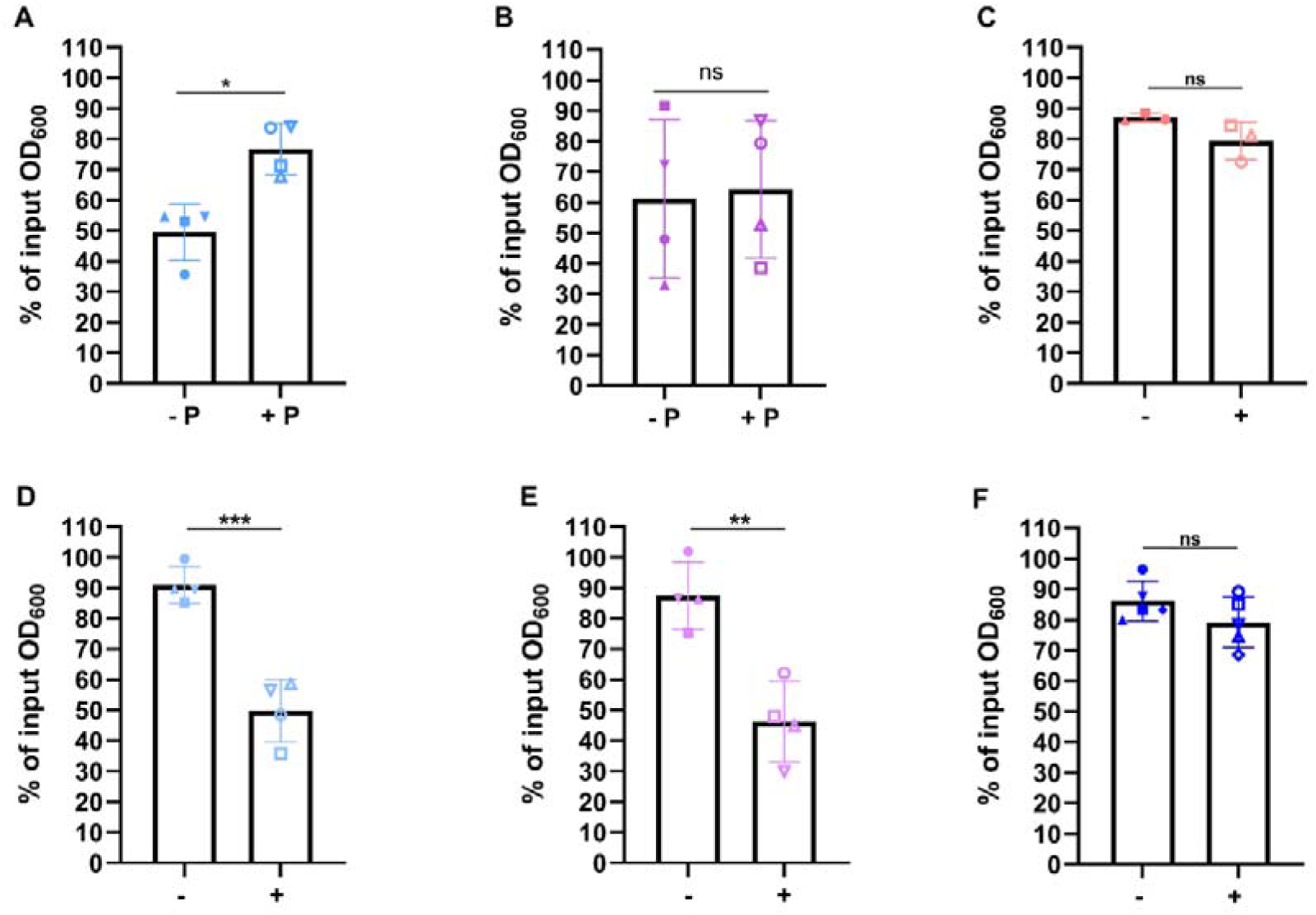
Plasmin cleavage disrupts ClfA-mediated clumping in fibrinogen. Stationary phase cultures of *S. aureus* ClfA^+^(A) and ClfA_ (B) were adjusted to OD_600_ = 1.5 in PBS and incubated in the absence and presence of plasmin (± P, 0.075 U/mL) for 1 h at 37°C. Bacteria were washed and incubated statically at 37°C for 105 min in the presence of fibrinogen (18.4 μg/mL). *S. aureus* ClfA^Δ40-214^ (C) ClfA^+^ (D) ClfA_ (E) and ClfA^−^ (F) were incubated in PBS with (+) and without (−) fibrinogen (18.4 μg/mL) for 105 min at 37°C. The OD_600_ after 105 min is expressed as a percentage of the original input OD_600_ at 0 h. Each symbol represents the % of input OD_600_ from a single matched experiment and the bars represent the mean value for each condition from four independent experiments. Error bars represent standard deviation, and statistical significance was determined by an unpaired student t-test. *, P < 0.05; **, P < 0.01; ***, P < 0.001; ns, non-significant.

Next the ability of *S. aureus* expressing a ClfA truncate that mimics the plasmin-cleaved state, (*S. aureus* ClfA^Δ40-214^), was tested for its ability to clump in a solution of fibrinogen (**Fig. 4C**). *S. aureus* ClfA^Δ40-214^ lost the ability to clump (**Fig. 4C**) while *S. aureus* ClfA^+^ (**Fig. 4D**) and *S. aureus* ClfA^R^ (**Fig. 4E**) incubated under the same conditions clumped in a solution of fibrinogen. *S. aureus* ClfA^−^ did not clump demonstrating that clumping activity is mediated solely by ClfA under the conditions used here (**Fig. 4F**). Together these results suggest that removal of residues 40-214 either by treatment with plasminogen or genetically, renders ClfA incapable of binding to soluble fibrinogen and promoting clumping of *S. aureus*.

### Truncation of ClfA enhances ClfA-mediated survival in human blood

To determine whether truncation of ClfA, and the resulting loss of fibrinogen binding, influences bacterial killing by the host in blood, the survival of *S. aureus* was monitored over three hours in freshly drawn human blood (**Fig. 5**). *S. aureus* ClfA^Δ40-214^ displayed a significantly increased survival in whole human blood compared to both *S. aureus* ClfA^+^ and the cleavage resistant variant ClfA^R^. There was no significant difference in the survival rates of *S. aureus* ClfA^+^ and ClfA^R^. Importantly, all strains grew similarly in serum (**Fig, S1**), indicating that the better survival of ClfA^Δ40-214^ compared to ClfA^+^ and ClfA^R^ is not due to a difference in the ability to grow in the liquid component of blood, but rather a difference in resistance to phagocytic killing. Together these results suggest that removal of the N1 domain increases ClfA-mediated survival in blood.

**FIG. 5.**
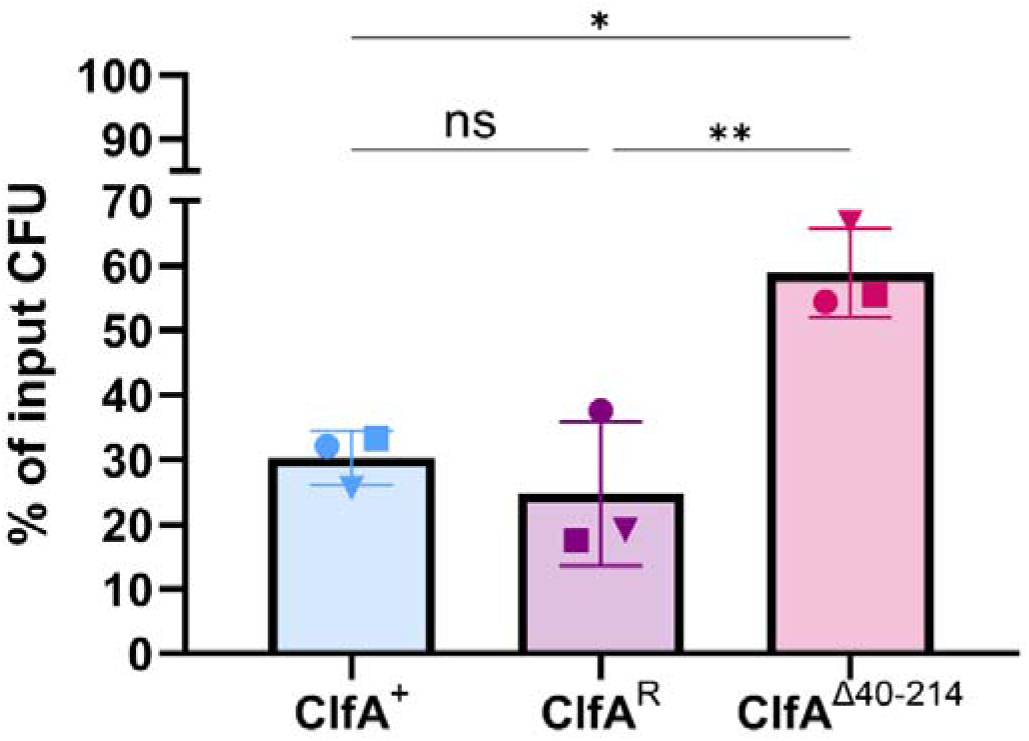
Truncation of ClfA enhances ClfA-mediated survival in human blood. Stationary phase cultures of *S. aureus* ClfA^+^, ClfAD and ClfA^Δ40-214^ were inoculated (ca. 2.5 × 10D CFU/mL) into freshly drawn human blood and incubated for 3 h at 37°C. Viable counts after 3 h are expressed as a percentage of the original input CFU at 0 h. Each symbol represents the % of input CFU recovered from a single experiment and the bars represent the mean value for each condition from three independent experiments. Error bars represent standard deviation, and statistical significance was determined by one-way Anova following Tukey’s multi-comparison test. Where significant P is represented: ns. P > 0.05; *, P < 0.05; **, P < 0.01.

### Truncation of ClfA increases kidney abscess formation by *S. aureus*

ClfA promotes renal abscess formation in murine infection models (4, 50–52). Having shown that the truncated form of ClfA interacts poorly with soluble fibrinogen (**Fig. 4**) but enhances whole blood survival (**Fig. 5**), we sought to determine how this might impact *S. aureus* infection in a mouse model. ClfA was not susceptible to cleavage by mouse plasmin in serum (**Fig. S2**) so the effect of truncation on the activity of ClfA *in vivo* was examined using the *S. aureus* expressing full-length ClfA and its variants. In agreement with our findings with human fibrinogen (**Fig. 4**), *S. aureus* ClfA^Δ40-214^ was impaired in its ability to induce clumping in a solution of mouse fibrinogen (**Fig. S2**). A sub-lethal intravenous dose of *S. aureus* ClfA^Δ40-214^ resulted in more severe abscess formation (**Fig**. **6A**) and a higher bacterial load (**Fig. 6B**) in the infected mouse kidneys compared to *S. aureus* ClfA^+^ or *S. aureus* ClfA^R^ indicating that the ability of *S. aureus* to disseminate to the kidney and establish renal abscesses is enhanced when residues 40-214 are missing from ClfA. There was no difference in bacterial load in the liver following infection with any of the strains (**Fig. 6C**). Comparing the survival of mice challenged with *S. aureus* expressing ClfA^+^, ClfA^R^ and ClfA^Δ40-214^ indicated that there were no significant differences in the survival rates of mice infected with any of the three strains over the 10 days of the experiment (**Fig. 6D**). The concentrations of IL-6 mice infected by all three strains was similar (**Fig. S3**). Together these data indicate that removal of residues 40-214 and therefore inactivation of the fibrinogen binding function of ClfA enhances the ability of *S. aureus* to form abscesses in mouse kidneys.

**FIG. 6.**
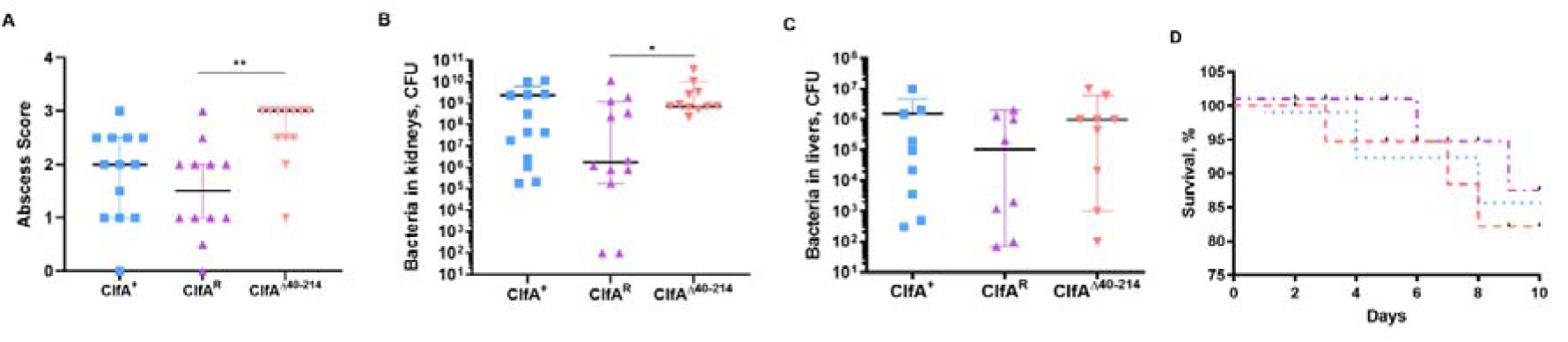
Truncation of ClfA increases kidney abscess formation by *S. aureus*. *S. aureus* ClfA^+^, ClfA^R^ and ClfA^Δ40-214^ were intravenously administered to NMRI mice (10 per group) via the tail vein with 200_μL of PBS containing 75 ng/mL anhydrotetracycline. Expression of ClfA was induced via an intraperitoneal injection of inducer anhydrotetracycline (5 µg/mouse) on Day 0, 1 h before infection and 1h post-infection. Kidneys and livers were collected aseptically, and kidney abscess scores (A) were assessed in a blinded manner; 0 represented healthy kidneys; 1 indicated one to two small abscesses without structural changes; 2 denoted more than two abscesses affecting less than 75% of the kidney tissue; and 3 indicated extensive abscess formation involving more than 75% of the tissue. Bacterial load in the kidney (B) and liver (C) were quantified by homogenising the organs and culturing on horse blood agar. (D) Mouse survival after infection. Percent survival of mice (n = 20 per group) after infection with 6 × 10^6^ CFU/mouse. The dashed line colour pairs to symbol colour for strains in panel A-C. Error bars represent 95% confidence intervals and statistical significance was determined by one-way Anova multiple comparisons with the Kruskal-Wallis post-test. *P < 0.05; **, P < 0.01. No symbol P > 0.05.

## Discussion

In this study we show that cleavage of ClfA by plasmin results in a pronounced loss of bacterial clumping in fibrinogen, reduced survival of *S. aureu*s in human blood and increased abscess formation in the kidney. We pinpoint the plasmin cleavage site to between residues R214 and A215 in ClfA (**Fig. 2**) and demonstrate that the activity of plasmin leads to the formation of a stable truncate of ClfA on the cell surface lacking residues 40-214.

*S. aureus* actively recruits plasminogen to its surface and converts it to plasmin using Sak (28), thus providing a mechanism to inactivate the fibrinogen-binding activity of ClfA on the cell surface, even in the absence of host-activated plasmin. Existing evidence supports fibrinogen binding to bacteria conferring protective benefits to the host (11, 13, 15, 51, 53–56) while conversely, other studies find it can drive pathogenesis (4, 12, 16, 49, 50, 52, 57–61). Such differences in outcome likely reflect infection model or strain-dependent factors. Here we used a sub-lethal intravenous challenge with *S. aureus* and followed survival of the infected animals and dissemination to the kidneys and liver. This showed that loss of fibrinogen binding in the truncate resulted in impaired bacterial killing, enhanced dissemination and increased abscess formation. This agrees with previous findings showing that mice expressing a fibrinogen truncate lacking the ClfA binding site demonstrate delayed bacterial killing and increased dissemination from the peritoneal cavity (16).

The proteolytic cleavage of cell wall-anchored proteins in *S. aureus* has been described previously. In some instances, surface-exposed cell wall-anchored proteins are completely degraded by proteases, such as FnBPA and FnBPB (62, 63). However, in other cases, cleavage at specific sites by bacterial or host proteases (24, 64, 65) serves as a post-translational modification that modulates protein function. For example, ClfB loses its ability to bind fibrinogen when cleaved by aureolysin (47) and SasG no longer inhibits bacterial clumping when processing by trypsin (25).

In this study, we demonstrate that removal of residues 40-214 from the mature ClfA protein abolishes its ability to interact with soluble fibrinogen. This finding was unexpected, as the canonical fibrinogen-binding domain of ClfA is thought to reside entirely within the N2 and N3 domains (residues 229-545) (18, 20). Interestingly, the loss of fibrinogen binding in the ClfA^Δ40-214^ variant mirrors the ligand-binding behaviour of ClfB, where cleavage by aureolysin outside of the minimal fibrinogen-binding domain results in a complete loss of function (47). These observations suggest that the ClfA-fibrinogen interaction is more complex than previously appreciated. While residues 229-545 are sufficient to support recombinant protein binding to a peptide derived from fibrinogen, they are clearly not sufficient to promote bacterial interactions with fibrinogen and fibrin in the bloodstream.

Elucidating the structure of the entire ClfA protein in complex with fibrinogen would reveal the full binding interface and uncover how ClfA engages fibrinogen/fibrin. This new finding highlights the drawbacks of studying host-pathogen interactions using recombinant bacterial adhesin domains *in vitro* and indicates that the interaction between ClfA and fibrinogen, and likely other MSCRAMMs and their ligands, is far more complex than the dock, lock and latch mechanism encapsulates and best studied using whole cells expressing the protein of interest.

Plasmin cleaves ClfA at the carboxyl terminus of arginine residues, consistent with its known substrate specificity (66). Proteolytic processing of ClfA results in the release of residues N-terminal of A215, yet the corresponding fragment (the majority of N1) is undetectable by SDS-PAGE suggesting it undergoes rapid secondary degradation. Recombinant N1N2N3 domains of MSCRAMMs often resolve into a single proteolytically stable fragment corresponding to N2N3, even in the absence of added protease (67). Degradation of the N1 domain of MSCRAMMs is likely driven by intrinsic disorder within N1 and trace protease contamination during recombinant protein preparation (25, 47, 67). Our *in vivo* experiments were conducted exclusively with a truncated ClfA variant as we found that murine plasmin was inefficient at cleaving ClfA (Fig. S2). This confirming that the observed phenotypes stem from loss of ClfA function rather than activity of the released fragment. However future studies should investigate if the released portion, or peptides derived from it, have biological activity.

## Funding information

M.B.T. was supported by a Postgraduate Ussher Fellowship Award from Trinity College Dublin. J.A.G. is supported by a Royal Society Wolfson Fellowship (Award 193002). Z.H. and T.J. are supported by Swedish Medical Research Council 2024-02667 and the Swedish state under the agreement between the Swedish Government and the county councils ALFGBG-1005076. E.C.L. and R.W.W. received support from Science Foundation Ireland Frontiers for the Future award 19/FFP/6484. R.W.W. received support from the Irish Research Council GOIPG/2019/4236. The funders were not involved with the design of this study, collection or analysis of data, decision to publish or preparation of the manuscript.

## Acknowledgements

We gratefully acknowledge the contribution to this study made by the University of Birmingham’s Human Biomaterials Resource Centre which has been supported through Birmingham Science City - Experimental Medicine Network of Excellence project. We thank Deirdre Muldowney and Kevin Lyons for technical assistance.

## Conflict of Interest

The authors declare there are no conflicts of interest of a financial or any other nature.

## Notes

### Competing Interest Statement

The authors have declared no competing interest.

### Summary of Updates

New figure added - figure 5 and supplemental data included

