## Supporting Information for "Plasmin-Mediated Processing of Clumping Factor A Impacts Bacterial, Aggregation and Abscess Formation during *Staphylococcus aureus* Infection"

**Table S1. Primers used in this study**

| **Primer** | **Description** | **Sequence** |
| --- | --- | --- |
| cSt_F | Inverse PCR for inclusion of C-terminal Strep-Tag II on pQE30::ClfA40-559 | *GCTGCTTGGTCTCATCCTCAATTTGAAAAATAAAAGCTTAATTAGCTGAGCTTG |
| cSt_R | Inverse PCR for inclusion of C-terminal Strep-Tag II on pQE30::ClfA40-559 | *CTCTGGAATTGGTTCAATTTC |
| cSt screen_F | Amplification of 166 bp region within cSt | ATAATTTGGCGCTCTATG |
| cSt_screen_R | Amplification of 166 bp region within cSt | ATCAACAGGAGTCCAAGCTC |
| cSt_SLIC_F | PCR to delete a 547 bp region of ClfA N1N2 from plasmid pCF-40 cSt | ACCCTGAAAATGTTAAAAAGACAGG |
| cSt_SLIC_R | PCR to delete a 547 bp region of ClfA N1N2 from plasmid pCF-40 cSt | GTTGAAACATTTTCCGCATTTGTA |
| ASAS_F | Amplification of a 547 bp region of ClfA N1N2 containing Ala_211_SerAlaSer_214_ mutation. | CTACAAATGCGGAAAATGTTTCAAC |
| ASAS_R | Amplification of a 547 bp region of ClfA N1N2 containing Ala_211_SerAlaSer_214_ mutation. | CCTGTCTTTTTAACATTTTCAGGGT |
| 214_F | Inverse PCR for removal of a 522 bp region of ClfA N1 on pQE30::ClfA40-559* | *GCATTTAGTTTAGCGGCAG |
| 214_R | Inverse PCR for removal of a 522 bp region of ClfA N1 on pQE30::ClfA40-559* | *TGCATCTGCTTCTTTACTGC |
| pQE30 MCS_F | Amplification of cloned sequence within pQE30 MCS. Screening primer. | GAGCGGATAACAATTATAATAG |
| pQE30_MCS_R | Amplification of cloned sequence within pQE30 MCS. Screening primer. | CCGAGCGTTCTGAACAAATC |
| pRMC2 MCS_F | Amplification of cloned sequence within pRMC2 MCS. Screening primer. | ATTCAGGCTGCGCAAC |
| pRMC2 MCS_R | Amplification of cloned sequence within pRMC2 MCS. Screening primer. | TTGTTGACATATATCATTG |

* = 5' phosphorylation; F = forward; R = reverse

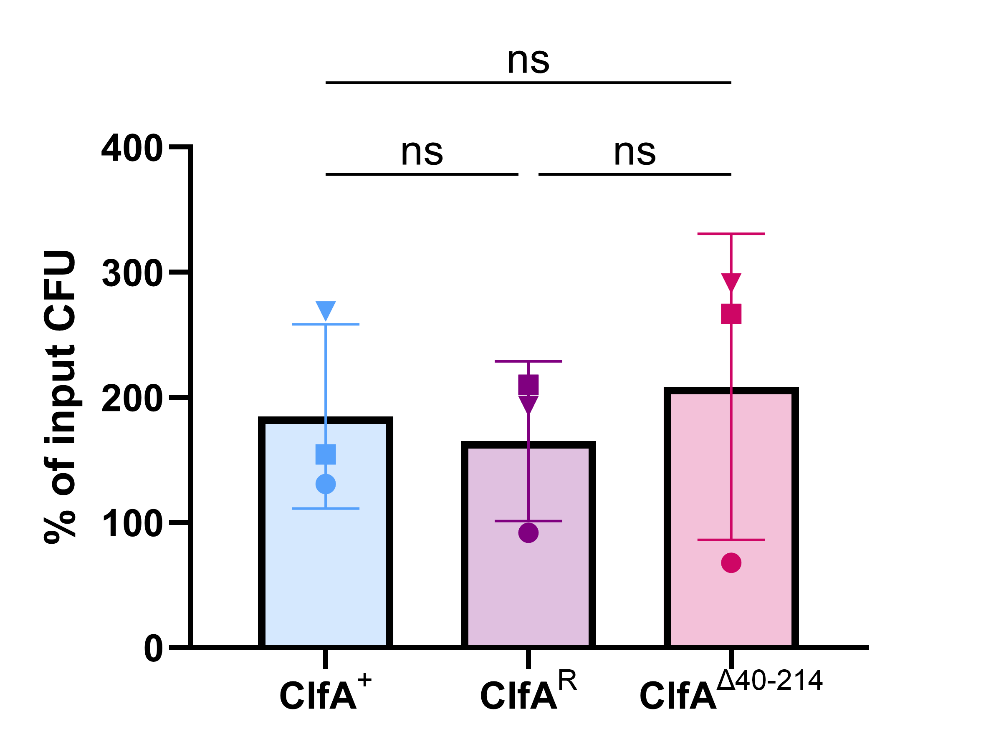

**FIG. S1. Truncation of ClfA does not alter *S. aureus* growth in human serum.** Stationary phase cultures of *S. aureus* ClfA^+^, ClfAᴿ and ClfA^Δ40-214^ were inoculated (ca. 2.5 x 10⁴ CFU/mL) into freshly drawn human blood and incubated for 3 h at 37°C. Viable counts after 3 h are expressed as a percentage of the original input CFU at 0 h. Each symbol represents the % of input CFU recovered from a single experiment and the bars represent the mean value for each condition from four independent experiments. Error bars represent standard deviation, and statistical significance was determined by one-way Anova following Tukey’s multi-comparison test. P > 0.05, no statistical difference (ns).

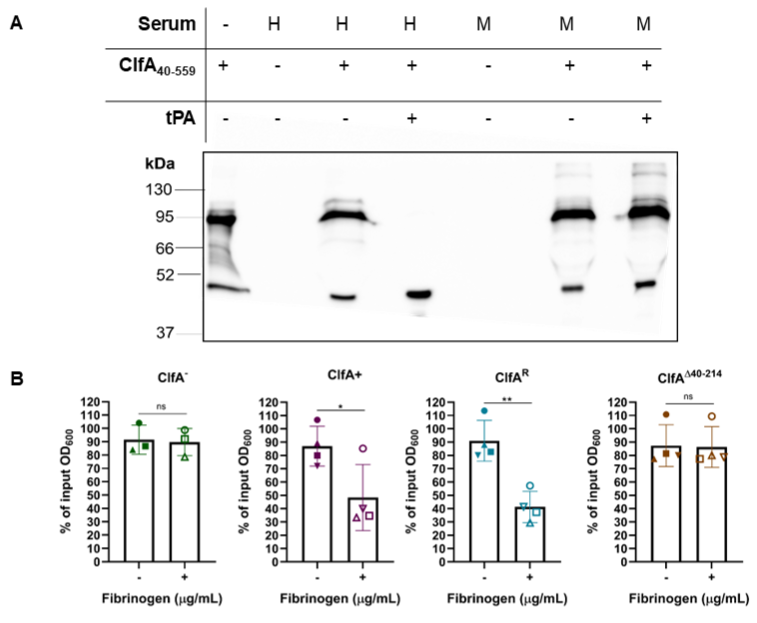

**FIG. S2. Cleavage of ClfA by murine plasmin and ClfA-dependent interactions of S. *aureus* with murine fibrinogen.** A) ClfA_40-559_ (5 μM) was incubated in human (H) or murine (M) serum (40%) that was pretreated with human tissue plasminogen activator (± tPA, 25 μg/mL) for 20 min at 37°C. Serum only controls did not include ClfA_40-559_. Samples were separated by SDS-PAGE and probed in a western immunoblot with anti-ClfA IgG (1:1000) and detected with protein A peroxidase (1:500). Blots represent two independent experiments. Interaction with soluble (B) murine fibrinogen. Stationary phase cultures of *S. aureus* ClfA^-^, ClfA^+^, ClfAᴿ, and *C*lfA^Δ40-214^, were adjusted to OD_600_ = 1 in PBS and incubated statically at 37°C for 105 min in the presence of M fibrinogen (18.4 μg/mL; C). Percent interaction was calculated by comparison to input OD_600_ at 0 h. Each symbol represents a single matched experiment. The bars represent the mean value for each condition from at least three independent experiments. Error bars indicate standard deviation, and statistical significance was determined by two-tailed unpaired student’s t-test . *, P < 0.05; **, P <0.01; P > 0.05, no statistical difference (ns).

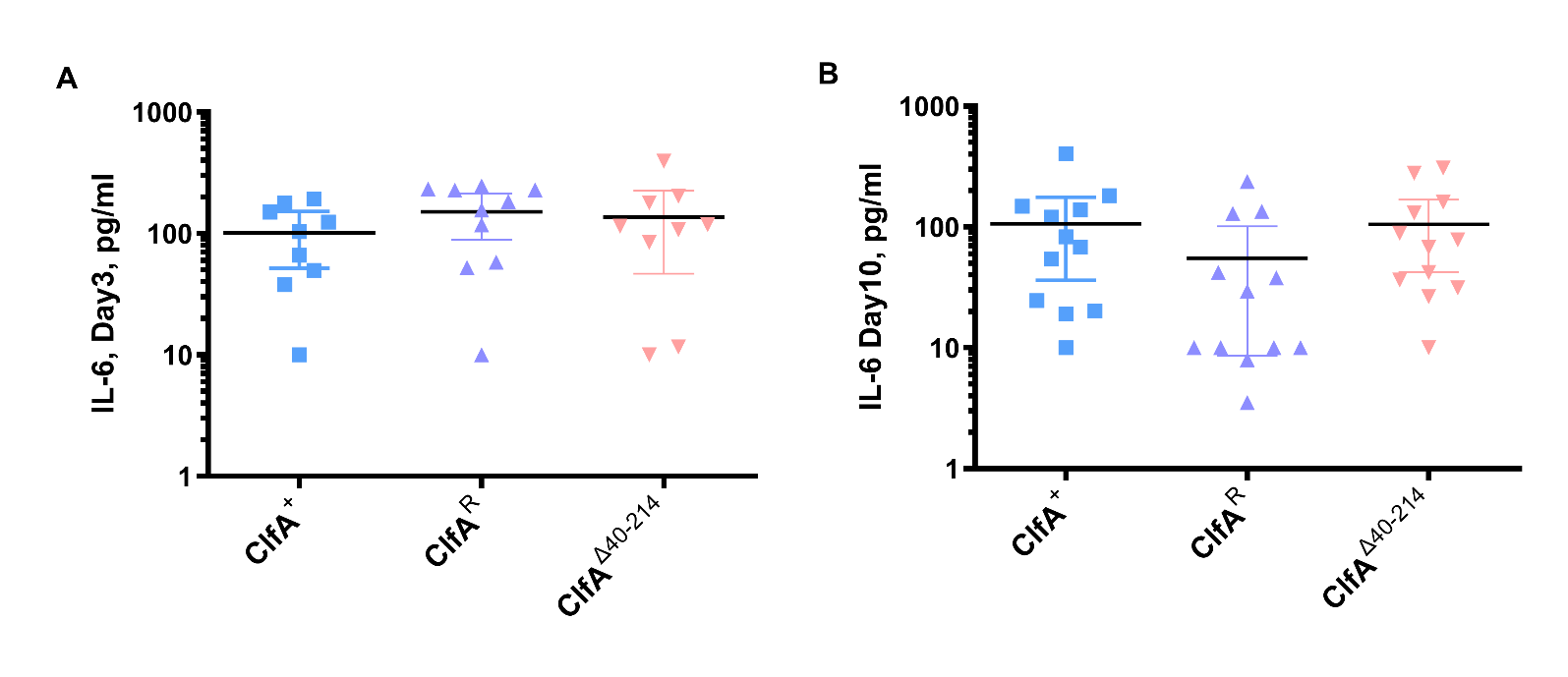
**FIG. S3. Truncation of ClfA does not induce a proinflammatory response.** *S. aureus* ClfA^+^, ClfA^R^ and ClfA^Δ40-214^ were intravenously administered into at least five mice per group for each experiment. Expression of ClfA was induced via a single injection of 200 microlitres of anhydrotetracycline (75 ng/mL) on Day 0, 1 h before infection. IL-6 in serum was measured on Day 3 (A) and Day 10 (B) by ELISA. Error bars represent 95% confidence intervals and statistical significance was determined by one-way Anova multiple comparisons with the Kruskal-Wallis posttest. P > 0.05, non-significant.
